# Spacemake: processing and analysis of large-scale spatial transcriptomics data

**DOI:** 10.1101/2021.11.07.467598

**Authors:** Tamas Ryszard Sztanka-Toth, Marvin Jens, Nikos Karaiskos, Nikolaus Rajewsky

## Abstract

Spatial sequencing methods increasingly gain popularity within RNA biology studies. State-of-the-art techniques can read mRNA expression levels from tissue sections and at the same time register information about the original locations of the molecules in the tissue. The resulting datasets are processed and analyzed by accompanying software which, however, is incompatible across inputs from different technologies. Here, we present spacemake, a modular, robust and scalable spatial transcriptomics pipeline built in snakemake and python. Spacemake is designed to handle all major spatial transcriptomics datasets and can be readily configured to run on other technologies. It can process and analyze several samples in parallel, even if they stem from different experimental methods. Spacemake’s unified framework enables reproducible data processing from raw sequencing data to automatically generated downstream analysis reports. Moreover, spacemake is built with a modular design and offers additional functionality such as sample merging, saturation analysis and analysis of long-reads as separate modules. Moreover, spacemake employs novoSpaRc to integrate spatial and single-cell transcriptomics data, resulting in increased gene counts for the spatial dataset. Spacemake is open-source, extendable and can be readily integrated with existing computational workflows.

## Introduction

Tremendous advances during the last decade led to high-throughput single-cell RNA sequencing technologies (scRNA-seq) that became the state-of-the-art for dissecting cellular heterogeneity within tissues. Spatial transcriptomics sequencing (STS) technologies present a further vital development that allows the assignment of single molecules to spatial positions, thus obtaining coordinates of gene expression. When spatial resolution is high enough to discern individual cells, this enables the identification of cell types and their interactions in spatial context. Spatial information is crucial in studying cell-cell communication mechanisms within the native tissue context and can yield new insights in disease states [1]. Recently published array-based methods are able to retain spatial information at different resolutions. Slide-seq (and Slide-seqV2) operates with 10μm beads that are evenly and randomly distributed on a 2D surface termed “puck” [2,3]. This size roughly corresponds to single-cell resolution. Other methods, such as spatial transcriptomics or the commercially available 10X Visium, work with a grid of 100μm diameter spots, regularly placed on a square glass (with 200μm distance between the centers), or 55μm diameter spots with 100μm distance between the centers, respectively [4,5]. These methods usually capture between 1-10 cells per spot, depending on the cellular density of the studied tissue. In more recent publications, high-definition spatial transcriptomics recovers gene expression at 2μm spatial resolution [6], while MiSeq Illumina flowcells were used to sequence mouse colon and liver tissues, achieving subcellular spatial resolution [7]. Fluorescent RNA labeling methods also achieve very high, often subcellular resolution, but operate on only a pre-selected panel of genes and are hence restricted to targeted studies of gene expression [1,8,9].

Akin to a technological revolution that took place with the advance of RNA-seq and scRNA-seq, we anticipate STS techniques to become invaluable for better understanding biological processes and mechanisms that lead to diseased states. Dissection of a tumor’s transcriptional heterogeneity is a prime example. Tumor progression is an intricate process that involves the coexistence of several cell types within the tumor, such as immune cells, native tissue cell types and abnormally growing tumor cells. While scRNA-seq can accurately identify different cell types and their transcriptional programmes, all spatial information regarding the cellular communication across cell types is lost. This information is critical to characterize spatial interactions within the tumor microenvironment and identify the mechanisms that create suitable conditions for the further progression of the disease, such as angiogenesis and hypoxia.

The various array-based STS methods differ not only in their experimental procedures, but also in the data they output and the associated software provided to process and analyze the raw data. Therefore, researchers who wish to take advantage of multiple methods need to get acquainted with several computational pipelines that operate with different logic and output structures. Such a situation can be time-consuming, perplexing, and can lead to the accumulation of errors when alternating between the different methods. There are a few computational processing tools available to date, namely the spaceranger from 10X [5], the ST pipeline [10] and slideseq-tools [2,3,10]. These tools, however, were developed for one specific STS technology (ST-pipeline and spaceranger for Visium and slideseq-tools for Slide-seq datasets), and are therefore not accommodating different types of data. Furthermore, they lack a unified framework to enable simultaneous processing of many different samples. Finally, they lack additional functionality, such as sub-sampling or merging of samples, integration of scRNA-seq with spatial datasets or support for troubleshooting of sequencing library construction by using long-read sequencing (Table 1).

**Table 1.** Comparison of spacemake with other published spatial-transcriptomics pipelines.

|  |  | slideseq-tools | spaceranger | ST pipeline | spacemake |
| --- | --- | --- | --- | --- | --- |
| **Input data** | Slide-seqV2 | ✓ | ✗ | ✓ | ✓ |
|  | 10X Visium | ✗ | ✓ | ✓ | ✓ |
|  | Seq-Scope | ✗ | ✗ | ✗ | ✓ |
|  | Other array-based STS | ✗ | ✗ | ✗ | ✓ |
|  | FISH | ✗ | ✗ | ✗ | ✗ |
| **Pipeline** | Customisable processing mode | ✗ | ✗ | ✗ | ✓ |
|  | H&E integration | ✗ | ✓ | ✗ | ✗ |
|  | Structured output | ✓ | ✓ | ✓ | ✓ |
|  | Parallel sample processing | ✗ | ✗ | ✗ | ✓ |
|  | Graphic QC reports | ✓ | ✓ | ✗ | ✓ |
|  | Automated downstream analysis | ✗ | ✓ | ✗ | ✓ |
| **Additional modules** | Saturation analysis | ✓ | ✓ | ✓ | ✓ |
|  | Technical replicate merging | ✗ | ✗ | ✗ | ✓ |
|  | novoSpaRc integration | ✗ | ✗ | ✗ | ✓ |
|  | Pacbio reads for troubleshooting | ✗ | ✗ | ✗ | ✓ |
| **General aspects** | Open-source | ✓ | ✗ | ✓ | ✓ |
|  | Extendable | ✓ | ✗ | ✓ | ✓ |

Here, we present spacemake, a unified computational framework for analyzing spatial transcriptomics datasets produced with Visium, Slide-seq, Seq-scope or any other STS technology. Importantly, spacemake performs data processing and downstream analysis in the same way, resulting in uniform reports and quality metrics that are easier to compare and interpret across different technologies. This renders spacemake an excellent candidate for multi-method projects. Apart from the standardized processing of raw data, spacemake can perform additional analyzes which we organize in different modules: downsampling and saturation analysis, merging of biological replicates, spatial reconstruction of scRNA-seq data or merging of scRNA-seq and STS datasets by using novoSpaRc [11] and analysis of long-read sequencing data for troubleshooting. Spacemake is written in snakemake [12] with a back-end logic written in Python. It provides an-easy-to-use command-line interface, through which it can be configured and run using a handful of commands. It readily works with various types of array-based STS methods and allows diverse, user-definable processing modes. Spacemake is versatile and can be used either as a new workflow, or be readily integrated into existing pipelines. Finally, spacemake is open-source and freely distributed through a Github repository.

## Results

### Spacemake processes different input data in a single workflow

Spacemake can handle different sequencing-based spatial-transcriptomic datasets, such as those stemming from - but not limited to - Slide-seqV2, 10x Visium or Seq-scope. In particular, it processes raw data (Illumina basecalls or fastq files) in identical fashion, regardless of the sequencing technology or the barcoding strategy of the spatial unit. As STS methods differ experimentally, we employ throughout the text the term spatial unit to describe the fundamental barcoded unit in space, e.g. beads, spots or clusters.

To allow for maximum flexibility, in spacemake each sample is associated with a set of ‘sample variables’, namely: a ‘barcode-flavor’, at least one ‘run-mode’, a ‘puck’ and a ‘species’ (Methods). The ‘barcode-flavor’ describes the barcoding strategy, that is, how the spatial unit barcodes and the unique molecular identifiers (UMIs) should be extracted from Read1 and Read2. The ‘run-mode’ parameter contains several variables which describe how the sample will be processed downstream and currently include: poly(A) and adapter trimming, tissue detection, multi-mapping read counting, intronic read counting, barcode cleaning, meshgrid creation and UMI cutoff (Methods). The ‘puck’ parameter allows the user to specify the spatial dimensions and bead diameter size of the underlying STS assay. Lastly, ‘species’ is a pair of a genome fasta file and an annotation file, from which spacemake will generate indices to be used later during mapping. After spacemake is configured and all parameters are set for all samples, it can be run, producing a unified and structured output for each sample (Sup. Fig. 1, Methods).

### Overview of the spacemake pipeline

Spacemake processes each sample starting from raw reads, which can be either Illumina basecalls, or demultiplexed fastq files. In the first case, spacemake demultiplexes the data using Illumina’s bcl2fastq2 tool [13]. Once raw fastq files have been created, a custom preprocessing script creates an unmapped BAM file: from each Read1, Read2 pair, a spatial unit barcode (or Cell Barcode, CB) and a UMI will be extracted and attached to the unmapped BAM file as CB and MI tags respectively. For each sample, this extraction is based on the previously defined barcode-flavor. Read sequences in this unmapped BAM come from Read2 sequences. Next, using Dropseq-tools [14] adapters and 3’ poly(A) stretches are optionally trimmed from each read. Reads are then mapped with STAR [15] and by using samtools [16] to input the unmapped BAM. After mapping, each read that maps to a gene body will be assigned a gene annotation using the TagReadWithGeneFunction command of Dropseq-tools. If the run-mode has multi-mapper counting turned on, spacemake will process the mapped BAM file line-by-line, and out of all possible alignments keep at most one alignment per read, to be counted later. Specifically, a multi-mapper is kept only if there is exactly one alignment to a genic region and all others to intergenic regions. In this case, the intergenic alignments are discarded. If a read aligns to multiple genes it is discarded. Finally, the digital gene expression (DGE) matrix is created using the DigitalExpression command of Drop-seq tools, with spatial unit barcodes used as whitelist (Sup. Fig. 1). After the DGE matrix is created, each sample is automatically analyzed: data filtering and clustering is done with scanpy [17] and the resulting data is saved as an hdf5 file. At the last step, web-based reports are generated by using Rmarkdown [18] and knitr [19] (Methods).

### Spacemake produces unified quality-control (QC) reports

Spacemake assesses the quality of each sample with multiple metrics. The commonly used FastQC [20] tool is first optionally called to assess sequencing library quality by flagging repetitive sequences, adapter content, GC bias, nucleotide composition and basecall qualities among others. Then, each sample is mapped to rRNA with bowtie2 [21] to assess the efficacy of poly(A) mRNA capture relative to abundant, contaminating ribosomal RNAs. After these QC steps are run, a per-sample web-based QC report is generated (Fig. 1). In particular, the number of genes, reads, UMIs and the reads/UMIs ratio are shown both as a histogram over all barcodes (Fig. 1A) and in tissue space (Fig. 1B). Randomness underlies the combinatorial complexity of the barcodes and is required for collision-free encoding spatial information. To assess the barcode randomness, a per-position nucleotide ratio is plotted, separated into quartiles by read counts, together with the Shannon entropy and the string compression length of the barcodes (Fig. 1C and 1D, Methods).

**Figure 1.**
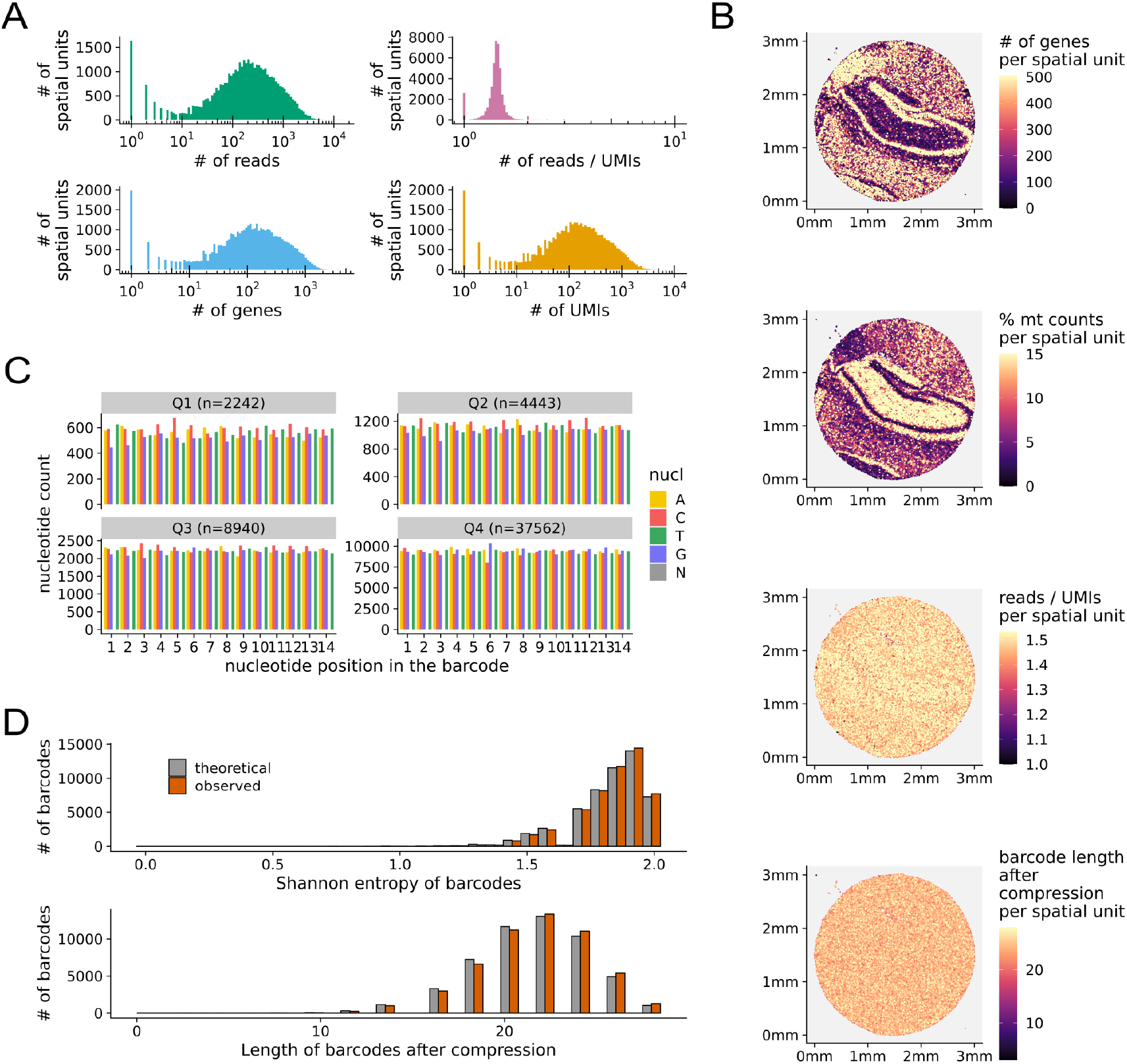
Spacemake produces uniform quality-control reports. **(A)** Histograms showing the number of genes, reads, UMIs and reads/UMIs ratio per spatial unit. **(B)** Quality control metrics plotted in tissue space. Top to bottom: number of genes, percentage of mitochondrial counts, reads/UMIs ratios and Shannon entropy, all shown per spatial unit. **(C)** Nucleotide frequencies per barcode position and quantile (segregated by the number of reads). **(D)** Shannon entropy and string compression length of the sequenced barcodes versus the expected theoretical distributions.

### Spacemake can readily aggregate spatial units

In some cases, it is useful to join nearby spatial units, effectively trading spatial resolution for statistical power by accumulating read counts (Fig. 2, Methods). This is particularly suitable for irregularly-spaced data points, such as Slide-seq, or when the data stems from an STS assay with subcellular resolution and is hence sparse, such as Seq-scope[7]. In addition, this aggregation also facilitates the comparison of spatial technologies operating at different resolutions, for instance Slide-seq and Visium.

**Figure 2.**
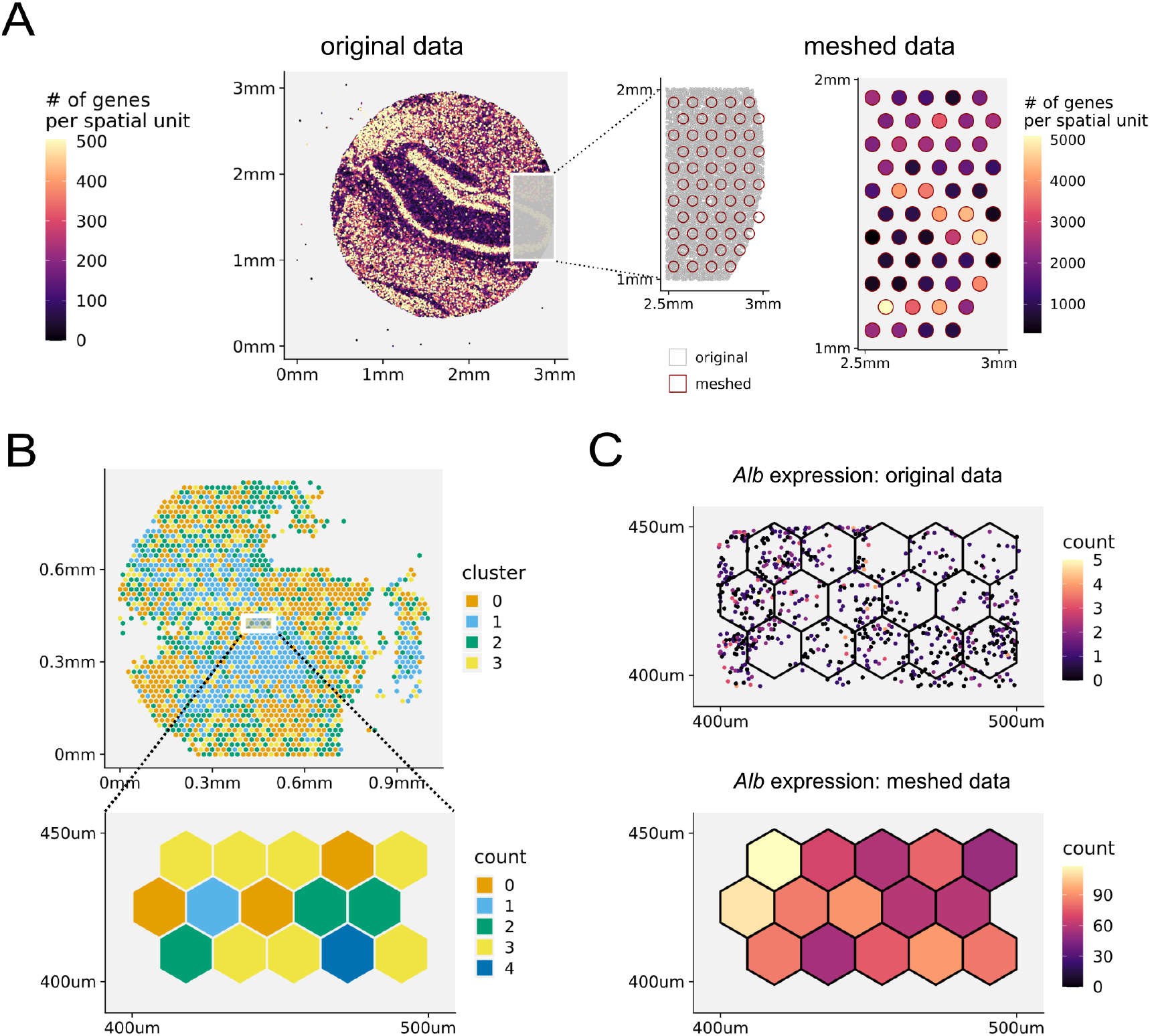
Spacemake seamlessly aggregates spatial units. **(A)** Spacemake can automatically create a Visium-style mesh grid (55μm diameters in a 100μm distance; also user-defined) and further processes the data mapped on this mesh. **(B)** Running on subcellular resolution datasets, such as Seq-scope, spacemake utilizes mesh-creation to join sub-cellular diameter spots into a 10μm-side hexagonal mesh. After the hexagonal mesh is created, downstream analyzes use it as input, *e.g* for cell type identification. **(C)** The highest expressed gene for this adult mouse liver sample is shown. Top right: raw-counts in the subcellular spots; bottom-right: counts assigned to hexagonal mesh cells.

In Seq-scope, for instance, ~800,000 barcodes spread out on a 1×1mm^2^ surface, so that the underlying diameter of each spatial unit is smaller than 1μm and contains a very low (not more than a few dozen) number of transcripts. To efficiently analyze such a sparse dataset, it is practical to create a ‘meshed’ grid (meshgrid) *in-silico*, where the diameter of each newly created spatial unit is 10μm, the approximate size of a eukaryotic cell. Spacemake offers two types of meshgrids out of the box: (1) a Visium-style meshgrid, where circles with a certain diameter are placed at equal distances from each other in a hexagonal grid (Fig. 2A); (2) a hexagonal meshgrid, where equal hexagons are created on top of the whole dataset, without holes in between (Fig. 2B). As the hexagonal meshgrid covers the entire area, no counts are discarded. For both meshgrids, spatial units falling into the same hexagon/circle are joined together and their gene expression counts are summed up (Fig. 2A,B).

### Downsampling analysis reveals library complexity and depth saturation

To assess library complexity and if saturation has been reached in scRNA-seq or STS experiments, a downsampling analysis is employed to estimate whether resequencing would result in a higher number of molecular counts per spatial unit. In spacemake, saturation analysis is implemented as a separate module (Fig. 3, Methods). First, the final BAM file is subsampled to 10%, 20%, …, 90% of the total reads using sambamba [22], and for each ratio a separate DGE matrix is generated. A saturation report is then compiled where median metrics are plotted as a function of the downsampling ratio (Fig. 3A). From the linearity of this curve it can be deduced that saturation has not yet been reached for this Seq-scope sample, even at 109 sequenced reads. In addition to plotting the median values, spacemake also reports histograms for each downsampling ratio per spatial unit, showing the global pattern rather than a single value per ratio (Fig. 3B, Sup. Fig. 4D).

**Figure 3.**
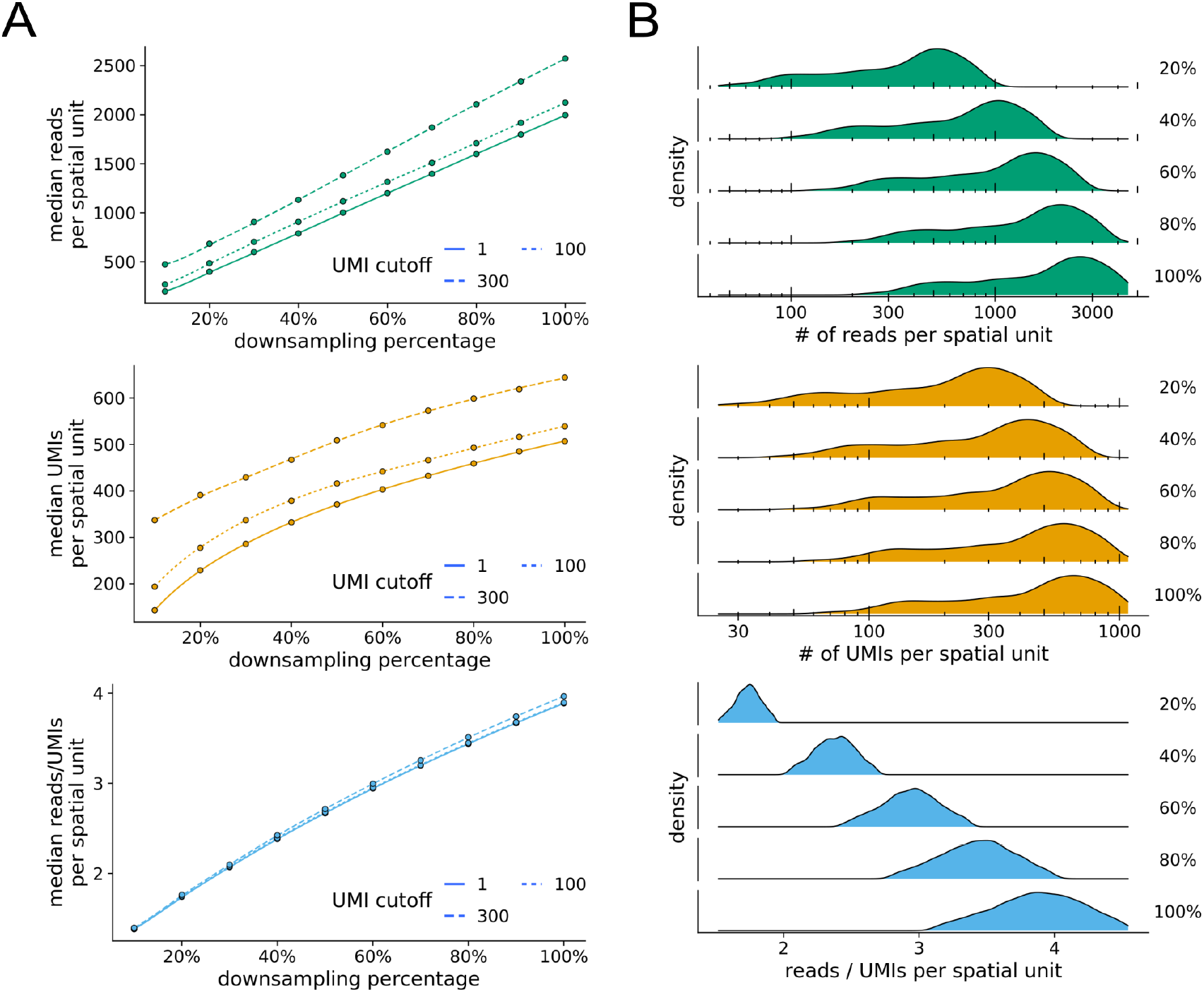
Spacemake can readily downsample the data to perform a saturation analysis. **(A)** Median number of reads, UMIs and reads/UMIs ratios per spatial unit are plotted against the downsampling percentage. Saturation analysis reveals that this Slide-seq sample hasn’t reached saturation yet, as the median UMIs curve hasn’t reached a plateau. **(B)** Density plots of a Seq-scope downsampled dataset.

### Spacemake can readily merge resequenced samples

Resequencing a library of sufficient complexity is a common practice to achieve higher molecular counts. In these cases the original and resequenced dataset have to be joined together, so that counts are quantified in the DGE matrix by properly removing duplicate reads. In spacemake, this process is implemented in the sample merging module which inputs the two separate, already processed datasets and joins them. Merging takes place at the level of the mapped BAM files. The resulting merged BAM file is further processed downstream, placing files in a new directory structure as if it was a separate sample.

### Spacemake offers a spatial reconstruction baseline of scRNA-seq data

Although spacemake is primarily designed to process STS datasets, it can also efficiently process data produced by the more standardized and popular scRNA-seq technologies. By now several pipelines exist for analyzing scRNA-seq data, for instance [23]. None of these, however, aims at incorporating a spatial reconstruction to the analysis. For this, spacemake utilizes novoSpaRc, a computational framework that reconstructs spatial information solely from scRNA-seq data based on the hypothesis that cells which are spatially neighboring also share similar transcriptional profiles [11,24]. Although novoSpaRc greatly benefits when a reference atlas of gene expression is available, its *de novo* mode is powerful and can yield insights into sub-structures of complex tissues, such as liver lobules, the intestinal epithelium or the kidney [24]. Spacemake employs novoSpaRc to yield a basic spatial reconstruction of scRNA-seq data that can serve as a baseline and can be used to derive further insights (Fig. 4, Table 2, Methods). Applied to a dataset of an adult mouse brain, for instance, spacemake recovers the basic structure representation of the mouse brain cortex (Fig. 4A).

**Table 2.** NovoSpaRc modes offered by spacemake and their outcome based on data availability.

| Data produced | Publicly available data | novoSpaRc mode | Outcome |
| --- | --- | --- | --- |
| - | scRNA-seq | *de novo* | basic spatial reconstruction |
| - | scRNA-seq + spatial | with markers | enhanced gene counts |
| scRNA-seq | - | *de novo* | basic spatial reconstruction |
| scRNA-seq | spatial | with markers | enhanced gene counts |
| spatial | scRNA-seq | with markers | enhanced gene counts |
| spatial + scRNA-seq | - | with markers | enhanced gene counts |

### Spacemake can integrate scRNA-seq data to a spatial transcriptomics dataset

When both spatial and scRNA-seq datasets of the investigated tissue are available, spacemake leverages novoSpaRc to integrate them. For this, the spatial dataset is regarded as a reference atlas and the scRNA-seq transcriptomes are mapped onto the locations of the spatial units. The integration of the two datasets leads to increased gene counts per spatial unit for the spatial dataset (Fig. 4B). Importantly, spacemake is not restricted to a specific technology but can utilize any spatial dataset as a reference atlas guiding the reconstruction. This becomes especially useful for widely studied or stereotypical tissues for which spatial datasets are already available, such as the adult mouse brain [25]. Mapping a publicly available scRNA-seq dataset [25] onto an existing spatial dataset [26], for instance, results in an enhanced number of genes per spatial unit (Fig. 4B). Furthermore, we observed that the expression profiles of spatially informative genes (identified with Moran’s I algorithm using squidpy’s spatial_autocorr function) become more distinct and more defined in space after novoSpaRc integration (Fig. 4C). To quantify the number of genes that are expressed in Visium spots after novoSpaRc integration, we modelled the expression of each gene by using a Gaussian Mixture Model with 2 components (Fig. 4D). Assuming that the lower (upper) mode of the bi-modal distribution describes low-to-no (low-to-high) expression, we calculated for each spot the number of genes expressed and compared it to the original data (Fig. 4B,D).

**Figure 4.**
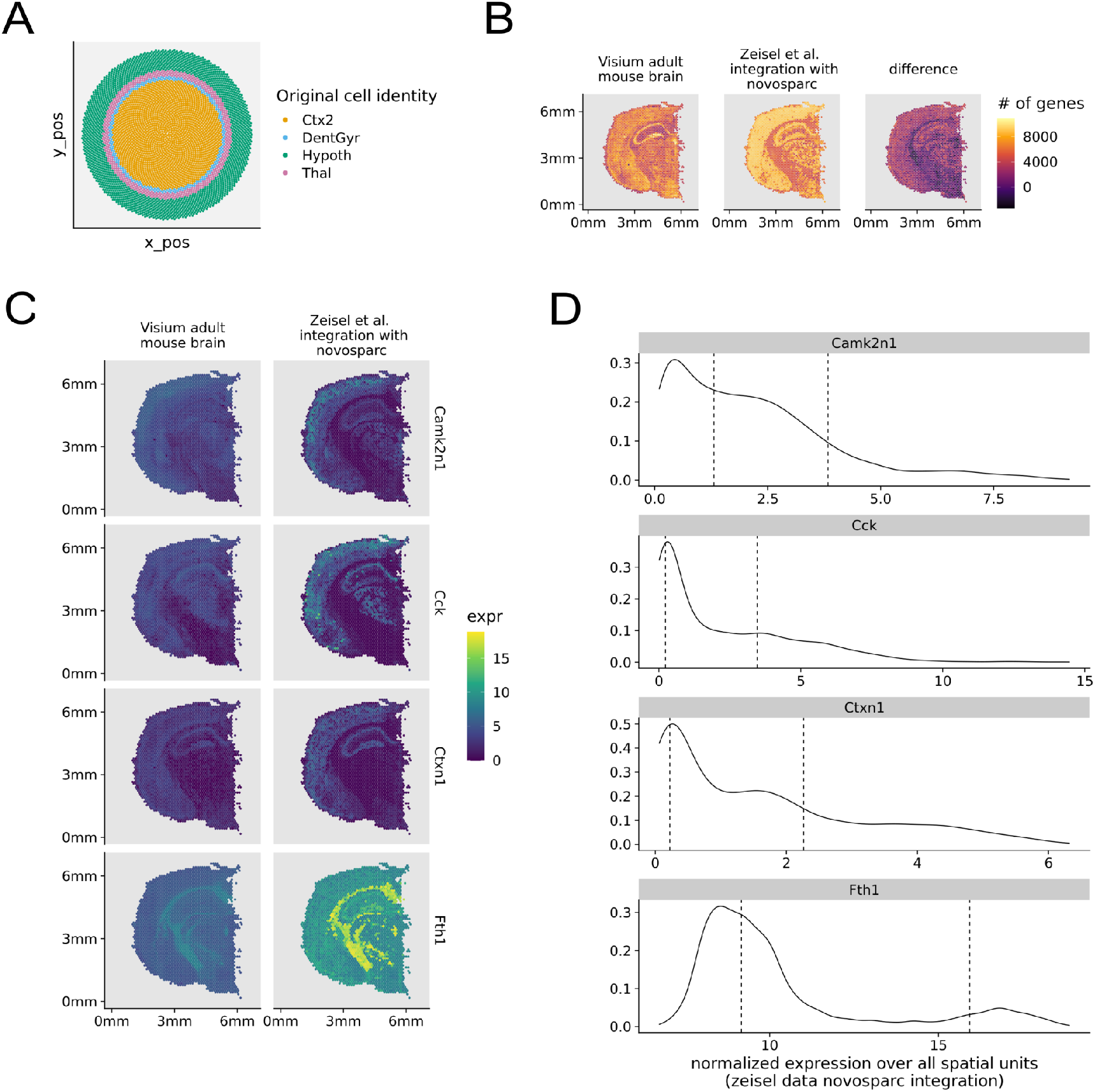
Spacemake can integrate scRNA-seq and spatial transcriptomics datasets. **(A)** Mapping an adult mouse brain scRNA-seq dataset with 30,000 cells onto an *in silico* created circular puck with 5000 locations reveals cortical layers. Tissue labels used: Thal, CA1, Hypoth, Ctx2, DentGyr, SScortex. **(B)** Integrating the single-cell and spatial transcriptomics datasets increases the number of genes quantified per spatial unit. **(C)** Expression of spatially informative genes as identified using squidpy. NovoSpaRc integration (right column) results in smoother expression patterns compared to the original ones (left column). **(D)** The bimodal distributions of gene expression are shown together with the corresponding mean values. To arrive at the results of panel (B), the expression of each gene was modeled with a Gaussian Mixture model with 2 components. For each spatial unit, only genes whose expression was in the upper mode were counted.

**Figure 5.**
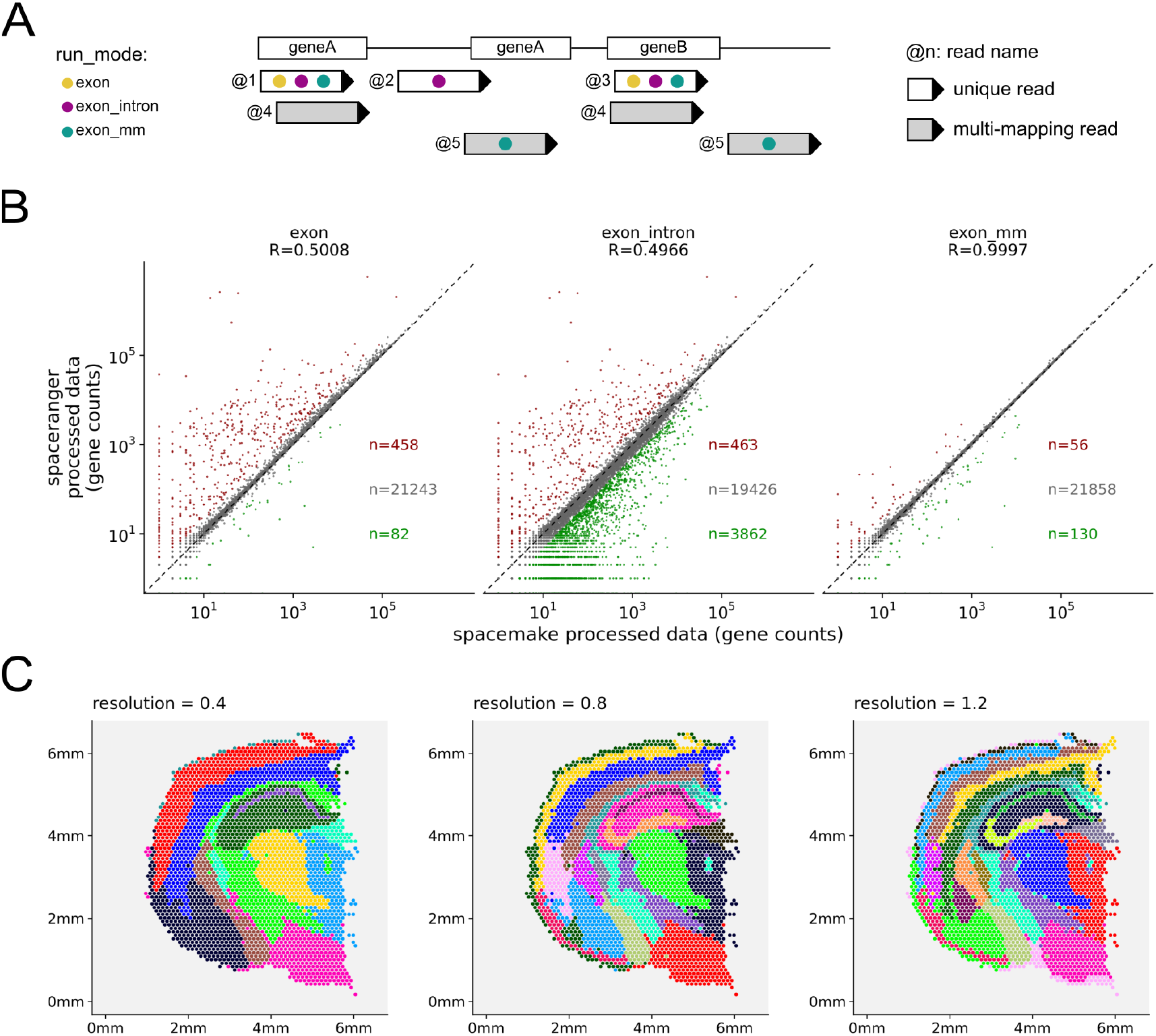
Spacemake offers several processing modes and produces a unified downstream output. **(A)** Spacemake can be run using several user-defined settings. Gene quantification depends on the run mode set to include reads mapping only on exons; on both exons and introns; on exons and intergenic regions; and whether the reads should be trimmed for poly(A)-tails and adapters. **(B)** Comparison of spacemake run-modes with spaceranger. Different run modes result in different. quantification with the highest correlation exhibited when only exonic reads are counted, multi-mappers are included and no trimming is performed. **(C)** For each sample and run-mode, clustering will be done on several resolutions. As we can appreciate, higher resolutions lead to more definable structures in space, at resolution we see a separation in the cortical layers as well as CA1/CA2, CA3 and the dentate gyrus

### Spacemake can leverage long-reads to troubleshoot library construction

Generation of STS and scRNA-seq libraries can be challenging due to the low amounts of RNA that may be captured from some samples. Especially when protocols are customized to accommodate specific experimental goals and needs, we have found it helpful to investigate our sequencing libraries by long-read sequencing. To this end, spacemake features a module to automatically annotate tens of thousands of long reads against a user-provided reference of expected adapter sequences and other oligo-nucleotides such as primers used during library construction (Sup. Fig. 3, Methods). The module then groups these annotations into recurring patterns of how these building blocks are arranged and provides an overview of the relative contributions of each class of such arrangements to the library. This allows the user to monitor cDNA integrity, for example from 10X Chromium beads (Sup. Fig. 3B,C), and enables to detect and subsequently mitigate potential primer and TSO concatenations as described in [27].

### Spacemake has flexible run-mode settings

A major strength of spacemake are the user-defined run-mode settings. A run-mode is created with the configuration command and provides complete control over how samples using this run-mode should be processed downstream (Methods). Adapter- and poly(A)-trimming can be turned on or off, and multi-mapper and intronic-read counting rules can be set. As each of these settings produces different results (Fig. 4A,B), it is often beneficial to initially run the analysis with several run-modes in parallel and then identify robust and reproducible results.

To demonstrate spacemake’s flexibility, we compared it against spaceranger on a publicly available adult mouse brain dataset [26]. We found that spacemake produced identical results to spaceranger when poly(A) trimming is turned off, only exonic reads are counted and multi-mapping read counting is turned on (Fig. 4A and Sup. Fig. 2).

Building on top of this flexibility in preprocessing and mapping, spacemake also allows to cluster the data using different parameters and saves all clustering results in the same automated analysis report (Fig. 4B). For the aforementioned dataset, higher clustering resolution leads to more biologically meaningful regions identified: at resolution 1.2, for instance, the pyramidal layer of the hippocampus separates into CA1/2, CA3 and the dentate gyrus (Fig. 4C).

### Spacemake provides automated downstream analysis

After processing is completed, spacemake performs a basic automated analysis of the data (Methods). For this spacemake employs scanpy [17] and squidpy [17,28]. More specifically, spacemake identifies cell types and their corresponding marker genes and plots them in an automatically generated report. If the user defines multiple UMI cutoffs for performing the downstream analysis, then multiple such reports are generated. For STS datasets in particular, spacemake uses squidpy to generate a cluster-to-cluster neighborhood enrichment heatmap (Sup. Fig. 4C), to calculate co-occurrence of spatial units and predict ligand-receptor interactions between spatial units.

We benchmarked spacemake against the results obtained in a Slide-seqV2 dataset [3]. For this, we first generated a raw fastq file from the slideseq-tools processed BAM file provided by the authors. Then, from the same file we created a DGE matrix using Dropseq-tools [14]. Finally, using the raw fastq files as input we ran spacemake and compared the results with the DGE matrix from the Slide-seqV2 BAM file. Spacemake achieves very high correlation with the Slide-seqV2 data, with most beads having a gene-gene correlation higher than 0.95 and the overall correlation being as high as 0.98 (Sup. Fig. 4B). Spacemake automatic clustering identifies spatially informative clusters, such as the cortical region, mouse hippocampus pyramidal layer, dentate gyrus and thalamic region, and the squidpy neighborhood enrichment analysis reveals spatial closeness of pyramidal-layer and cortical neurons (Sup. Fig. 4C).

### Spacemake is fast and scales with number of reads

Spacemake is fast, scalable and supports multithreaded processing. To benchmark spacemake, we processed the publicly available adult mouse 10X visium data using both spaceranger and spacemake. We observed that when using 6 cores, spacemake is 1h faster than spaceranger while producing the same results (Sup. Fig. 5A). Spacemake also scales well with the number of reads: for the Slide-seqV2 sample with 70 million reads, total run time was just over 1h, while 1 billion Seq-scope reads took 18 hours to process (Sup. Fig. 5A,B). Moreover, spacemake can run several samples in parallel. For a single sample, spacemake requires 4 cores minimum to run, so that with 8 or 12 cores several samples can be processed together, thus starkly reducing the average running time per sample.

## Discussion

As spatial sequencing technologies become increasingly available, the existence of robust, reproducible bioinformatics pipelines is of paramount importance. Here, we present spacemake, a comprehensive computational framework that efficiently analyzes spatial transcriptomics datasets stemming from different technologies. Spacemake is extendable, scalable and provides a complete solution from processing of raw data, over several quality controls and automated reports all the way to advanced downstream analyzes. Spacemake’s core strength is the unified processing of different data types, rendering it highly suitable for projects that use multiple methods. Spacemake is open-source, freely available and can be smoothly integrated with other packages that perform downstream analysis [28].

Spacemake is highly modular. It currently contains modules for downsampling and saturation analysis, sample merging, a baseline spatial reconstruction of scRNA-seq datasets and analysis of long-reads, and can be readily extended to add more functionality. Moreover, spacemake is versatile enough and can be used to analyze not only spatial transcriptomics datasets, but also scRNA-seq data. To demonstrate spacemake’s capabilities, we have used it to process and analyze Slide-seqV2 and 10X Visium datasets, showing that spacemake accurately reproduces the processed data of the two technologies. We further illustrated how spacemake can integrate scRNA-seq and STS datasets by employing novoSpaRc.

It should be noted that currently, spacemake processes and analyzes sequencing data, but not imaging data. Some spatial transcriptomics techniques, however, require to register the barcodes of the beads or spots in space by imaging. In a companion paper, some of us present a complete computational framework for efficiently handling such datasets, called Optocoder [32]. Spacemake can be readily integrated with Optocoder or similar methods.

Finally, it would be useful to extend spacemake to handle different types of data, e.g. protein expression or chromatin state. As novel techniques that provide diverse molecular readouts from the same cell are being constantly developed, it will be essential to possess a unified framework that can process the different data modalities. We plan to extend spacemake to accommodate such datasets in the future.

## Methods

### Run-mode settings

For each sample one or multiple ‘run-modes’ are defined to describe how spacemake should process it downstream. Each run-mode has a name and several parameters: automatic tissue detection (on/off), poly(A) and adapter trimming (on/off), intronic read counting (on/off), multi-mapping read counting (on/off), data meshing (on/off), number of expected barcodes, UMI-cutoff, DGE matrix cleaning (on/off). Each of these parameters are set through the command line. Currently, spacemake offers the following run-modes out of the box: scRNA_seq, visium, slide_seq and seq_scope, with parameters corresponding to each technology.

### Data preprocessing and mapping

The publicly available datasets were obtained as described in the data availability section below. FastQC (v0.11.9) was used to assess sequencing quality and a Python custom script was used to retrieve the cellular barcodes and UMIs for the different read structures (Visium: R1[1–16] for the spot barcode and R1[16–24] for the UMI and cellular barcodes; Seq-scope: R1[1–20] for the bead barcode and R2[1–9] for UMI; Slide-seq: R1[1–14] for bead barcode and R2[15–23] for UMI). During the barcode and UMI retrieval an unmapped BAM was created where each R2 sequence was tagged with the correct cell-barcode and UMI.

### Poly(A) and adapter trimming

If poly(A) and adapter trimming is switched on for the current run-mode, the 3’ ends of reads are trimmed for poly(A) and overlapping user-defined adapter stretches. This processing is performed with the functions TrimStartingSequence and PolyATrimmer of Drop-seq tools (v2.4.0) for poly(A) and adapter trimming respectively.

### Mapping and gene tagging

Alignment to the genome was performed with STAR (v2.7.9a) using the unmapped BAM as input and under the default parameters. The following genomes and annotation files were used: mm10 & M23 and were downloaded from Gencode. Gene tags were added with the function TagReadWithGeneFunction of Drop-seq tools.

### Multi-mapping read counting

Multi-mapping reads were counted using a custom python script which parsed the read-name sorted (STAR default output) final BAM line-by-line. For each read name, maximally one read was kept. If a read mapped to several genomic locations - but only one exonic region - this exonic-mapping read was kept and the rest were discarded. If a read mapped to several exonic locations it was removed altogether. During parsing, each kept read was flagged as primary, and the parsed output (now containing at most 1 read for each multi-mapper) was piped into the DigitalExpression of Dropseq-tools, which was run with a MAPQ=0 filter, to ensure multi-mapper inclusion.

### DGE creation

Once the aforementioned steps are run, the DGE matrix is generated. If the provided dataset contains a list of spatial barcodes, it is used as a ‘whitelist’. Otherwise, snakemake uses the n_beads parameter of the current run-mode to select the top n_beads number of barcodes with the highest read count using the BamTagHistogram function of Dropseq-tools. Finally, the DGE matrix is generated using either the ‘whitelist’ of spatial barcodes or the top n_beads barcodes.

### DGE barcode cleaning

For a user-defined set of primers, spacemake can optionally discard barcodes that overlap with any of these primers. This is controlled by the clean_dge parameter of a run-mode. When set to true, the following barcodes are removed: (1) barcodes that have at least 4nt overlap with any of the primers in the 3’-end; (2) barcodes that have an at least 7nt overlap with any of the primers, anywhere in the barcode itself. If selected, this step is run before generating the DGE matrix.

### Tissue detection

For the samples that was required, spacemake performed tissue detection as follows: first, for each spatial unit its neighboring spatial units were computed. For 10X Visium that is straightforward, as the data points lie within a hexagonal grid. For irregular grids such as Slide-seq datasets, we created a meshgrid and then quantified the spatial unit neighborhoods. This resulted in the generation of contiguous areas. The largest contiguous area was then considered to be under the tissue.

### Automated downstream analysis

For downstream analysis the text based DGE matrix was first parsed line-by-line using a custom python script to create a sparse matrix (Compressed Sparse Column), and cast as an AnnData object, and finally saved in h5 format to ensure minimal space. Then, the standard scanpy single-cell workflow followed with default parameters. We selected the top 2,000 highly variable genes and 40 principal components to use for clustering using the leiden algorithm [29] and lower-dimensional representation with UMAP [30]. Each sample was clustered using the scanpy.tl.leiden functions and for several resolution values. Cell type markers were identified with the rank_genes_groups function. For STS datasets squidpy was used by running the built-in squidpy.gr.spatial_neighbors function. Spatial co-occurrence was computed with squidpy.gr.co_occurrence and the ligand-receptor analysis with squidpy.gr.ligrec.

### Meshgrid creation

We created the mesh grids *in silico* using the numpy.mesh function. For both grids (Visium-style and hexagonal), a rectangular grid was first created with spot_distance_um (spacemake parameter - user definable) horizontal distances and sqrt(3) * spot_distance_um vertical distances. This mesh was then duplicated and spatially translated, so that the result of the two meshes was a mesh where the distance between any two neighboring points was exactly spot_distance_um. For the Visium-style mesh we joined beads which fall into any circle (with mesh points as circle centers) with a diameter of diameter spot_diameter_um. For the hexagonal mesh we calculated the distance between each spatial unit in the data and the mesh-points, and for each spatial unit we selected exactly one mesh point, the one with the minimum value.

### Downsampling analysis

Downsampling analysis was done by first splitting the final BAM file into different percentages with sambamba (v0.6.8). Then the downsampled BAM files were fed into the same processing pipeline described above for further analysis.

### Spatial reconstruction with novoSpaRc

The *de novo* spatial reconstruction of the adult mouse brain scRNA-seq data was done with novoSpaRc (v0.4.3) and by using the default parameters and a circular disk as a target space. The top 100 highly variable genes were selected for the reconstruction. For the spatial reconstruction with markers, the corresponding Visium dataset was used to first create a reference atlas. The top 200 highly variable genes were first obtained (both from Visium and single-cell datasets) and 195 of them remained after intersecting them. Reconstruction was done with novoSpaRc and with parameter alpha=0.5.

### Long-read analysis

The cDNA molecules should contain specific oligo-nucleotide building blocks in the right places, in addition to mRNA sequence and (parts of) the original poly(A) tail. Spacemake first aligns a catalog of such building blocks (SMART primer handles, poly(T), Template Switch Oligo, Illumina sequencing adapters, etc.) via local Smith & Waterman to each read. These alignments are then analyzed jointly for each long read and condensed into “signatures” which identify the presence/absence and relative ordering of each building block. Finally, the observed signatures are counted, compared systematically against the expected signature (for example: P5, bead_start, poly(T), N70X for a DropSeq bead-derived Illumina library) and the following diagnostic plots are generated: graphical breakdown of the library by signatures, zoom-in on bead-related features, mismatch and deletion analysis, as well as histograms of start/end positions for each part of the expected signature. We acquired the publicly available data as described below and every 250th read was selected and analyzed with the spacemake.longread module using the ‘chromium’ longread-signature.

### QC reports

QC plots were created with custom R scripts based on the ggplot2 package (v3.3.5). The automatically generated QC sheets were created with a custom Python script that collected the plots into a single pdf file. The Shannon entropy for each spatial unit barcode BC was calculated using the following formula:

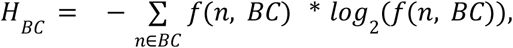

where *f*(*n*, *BC*)is the relative frequency of a nucleotide *n* in barcode *BC*. The length of string compression for a spatial unit barcode was calculated the following way: first the barcode BC was compressed (such that AAACATTA becomes 3A1C1A2T1A) and then the character-length of this compression representation was returned. The observed values were compared against theoretical values as follows: random barcodes were first generated for each sample and their Shannon entropy and string compression were then computed. The number of random barcodes generated was always the same as the number of real barcodes, for all samples.

### External DGE processing

Spacemake offers the possibility to process external count data. In this case, instead of starting from the raw data, the sample is processed downstream from the DGE matrix creation. Spacemake will perform the automated analysis and clustering and generate the corresponding reports.

## Code availability and requirements

Spacemake is freely available and can be found on Github: https://github.com/rajewsky-lab/spacemake

License: GPLv2

Operating system: Unix

Programming language: Python, R

Requirements: Python 3.6 or higher. R 4.0 or higher.

## Data availability

The Slide-seqV2 adult mouse brain dataset was downloaded from https://singlecell.broadinstitute.org/single_cell/study/SCP815/highly-sensitive-spatial-transcriptomics-at-near-cellular-resolution-with-slide-seqv2 (Puck_200115_08).

The 10X Visium dataset was downloaded from https://support.10xgenomics.com/spatial-gene-expression/datasets/1.0.0/V1_Adult_Mouse_Brain.

Seq-scope data was downloaded from https://www.ncbi.nlm.nih.gov/geo/query/acc.cgi?acc=GSE169706. For the analysis shown in this paper the dataset from healthy mouse liver with accession id SRR14082759 was used.

We used tile Nr. 2105 and extracted the bead barcodes and their positions were from raw fastq files found here https://deepblue.lib.umich.edu/data/concern/data_sets/9c67wn05f?locale=en, with the help of Seq-scope’s own script available here: https://github.com/leeju-umich/Cho_Xi_Seqscope/blob/main/script/extractCoord.sh

For the single-cell and novosparc mapping we used publicly available adult mouse brain data from [25], available here: https://storage.googleapis.com/linnarsson-lab-loom/l5_all.loom. We only used tissue labels comparable with the spatial Visium sample, namely: Thal, CA1, Hypoth, Ctx2, DentGyr, SScortex. We processed the data using spacemake and by treating them as an external DGE matrix.

For long-read sequencing data we used a subset of reads from SRR9008425 and SRR9008429, which were nanopore sequenced cDNA sequences derived from 10X Chromium beads from [31].

## Acknowledgements

N.K. was supported by DFG grant RA 838/5-1 and DFG grant KA 5006/1-1. T.R.Sz-T. acknowledges funding from the European Union’s Horizon 2020 research and innovation program, under the Marie Sklodowska-Curie Actions (MSCA) grant (721890) and funding from DFG Excellence Cluster 2049/Neurocure.

## Author contributions

N.K. conceived, designed and implemented the initial version of the pipeline. T.R.Sz.-T. implemented the pipeline in Snakemake. T.R.Sz.-T. and M.J. developed the pipeline. M.J. designed and implemented the long-read analysis module. T.R.Sz.-T. performed all computational and data analyses except for the long-read analysis which was performed by M.J. N.K. and N.R. supervised the study. All authors wrote the manuscript.

