## Supplementary figures for "Spacemake: processing and analysis of large-scale spatial transcriptomics data"

### Supplementary Data and Figures

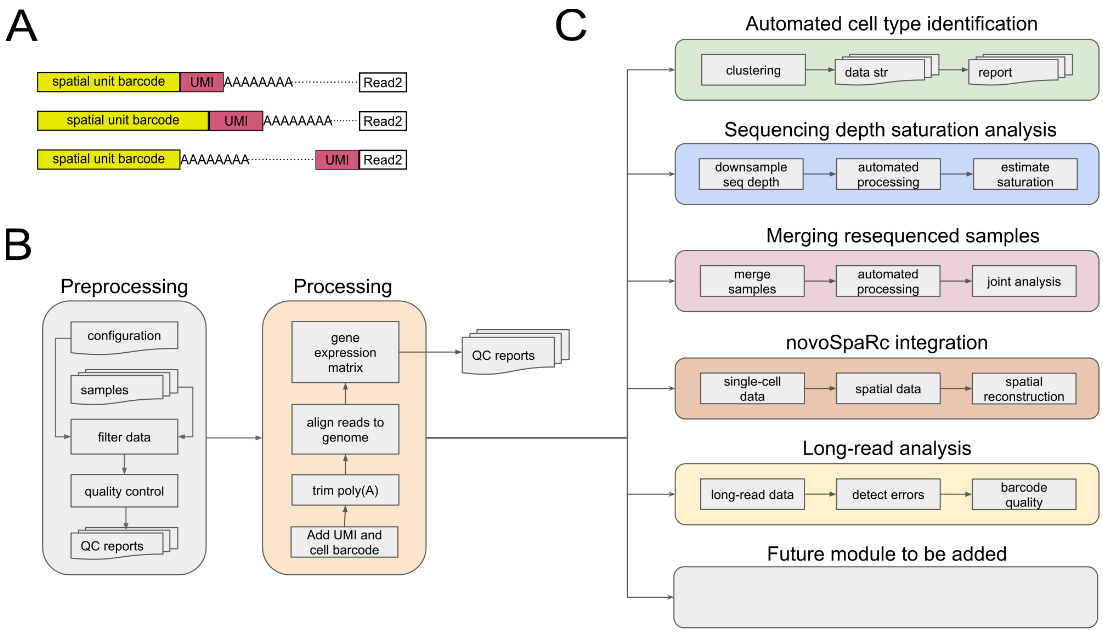


**Supplementary Figure 1.** Overview of spacemake. **(A)** Spacemake can handle various types of inputs: Read1 and Read2 cell-barcoding strategy is flexible. **(B)** Preprocessing, QC and processing steps. Each sample is processed the same way, regardless of the input type. **(C)** Spacemake is modular and extendable. Each module is implemented with a separate set of rules and commands, and everything is put together in the top level Snakefile.

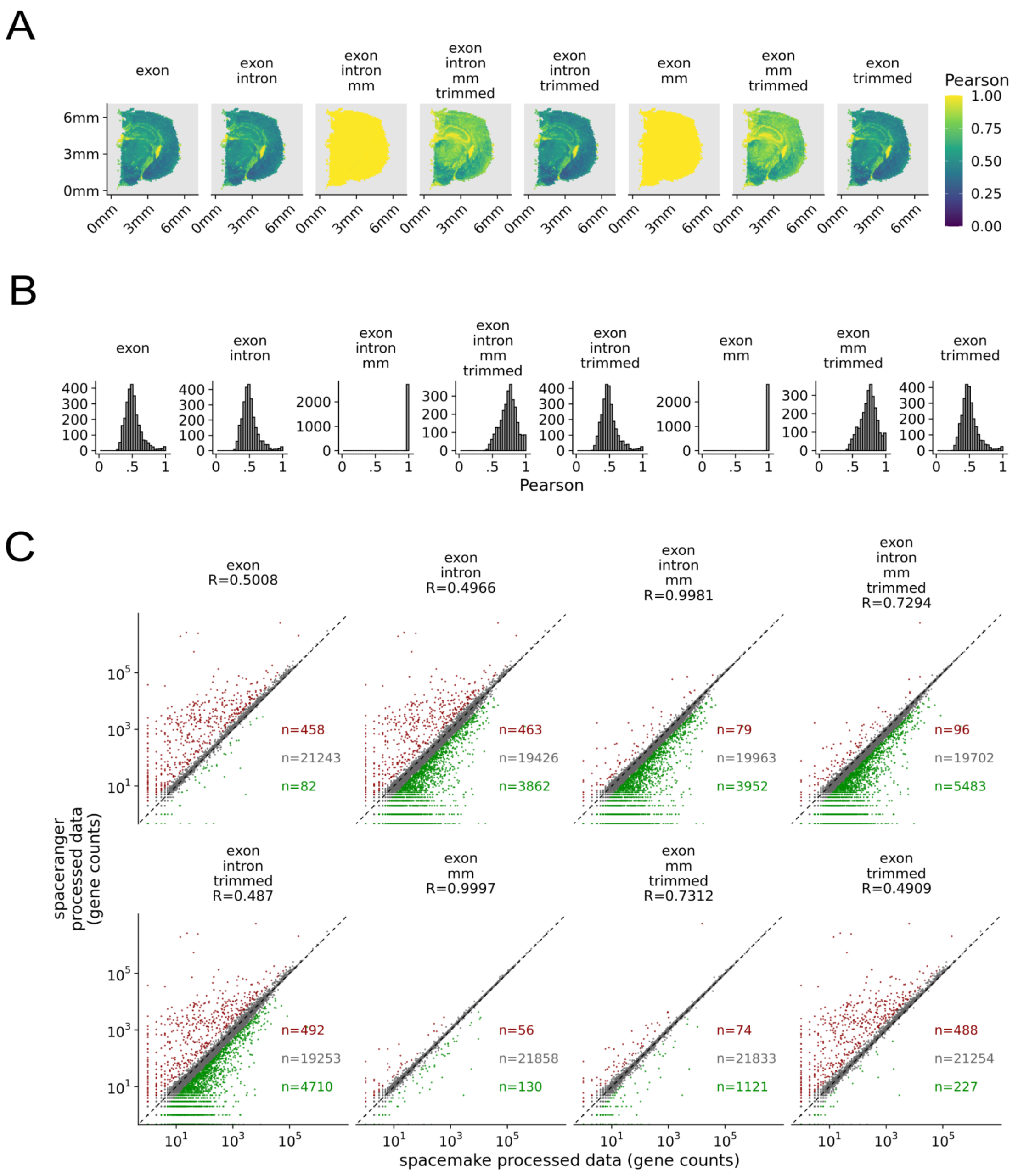


**Supplementary Figure 2.** Spacemake offers customizable run-mode settings and correlates well with spaceranger. **(A)** Correlations of spacemake run-modes with spaceranger. **(B)** Histograms of Pearson correlations for the different run-modes. **(C)** Correlations of aggregated gene counts. Each dot represents a gene. Red-colored genes exhibit an at least 2-fold increase in spaceranger vs spacemake, while green-colored genes exhibit an at least 2-fold increase in spacemake vs spaceranger.

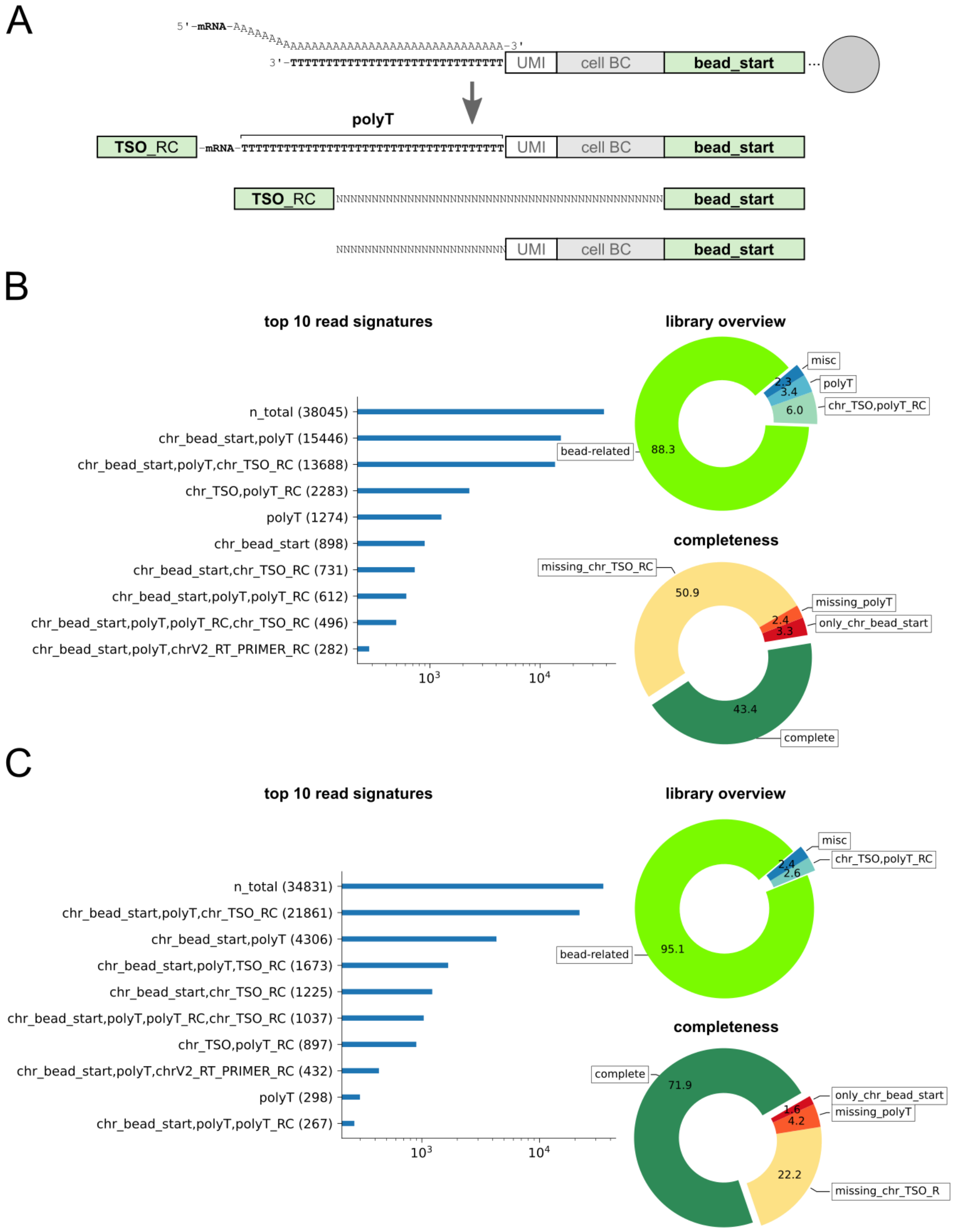


**Supplementary Figure 3.** Nanopore long-read analysis **(A)** Capture and reverse transcription of mRNA molecules with barcoded beads (top) can lead to the production of different cDNA species (bottom). The expected product contains Template Switch Oligo (TSO), mRNA, poly(T), and the bead-side primer handle labeled 'bead_start'. Identifiable building blocks labeled in bold-face. **(B, C)** Spacemake long-read module annotation of 10X Chromium-derived cDNA from [31]. Left: the most common signatures are plotted on a horizontal bar-chart (log-scaled x-axis). Right, top: overview donut-plot showing that the majority of cDNAs contain the expected bead primer-handle (bright green). Right, bottom: breakdown of the primer-handle containing cDNAs reveals that < 5% lack a detectable poly(T) tract and that 51% (B, SRR9008425) and 24% (C, SRR9008429) of reads are not terminated by identifiable TSO sequences. Note that oligo block labels for (B), (C) include ‘chr_’ as a prefix for 10X Chromium specific sequences, whereas (A) applies more broadly, for example also to Drop-seq beads.

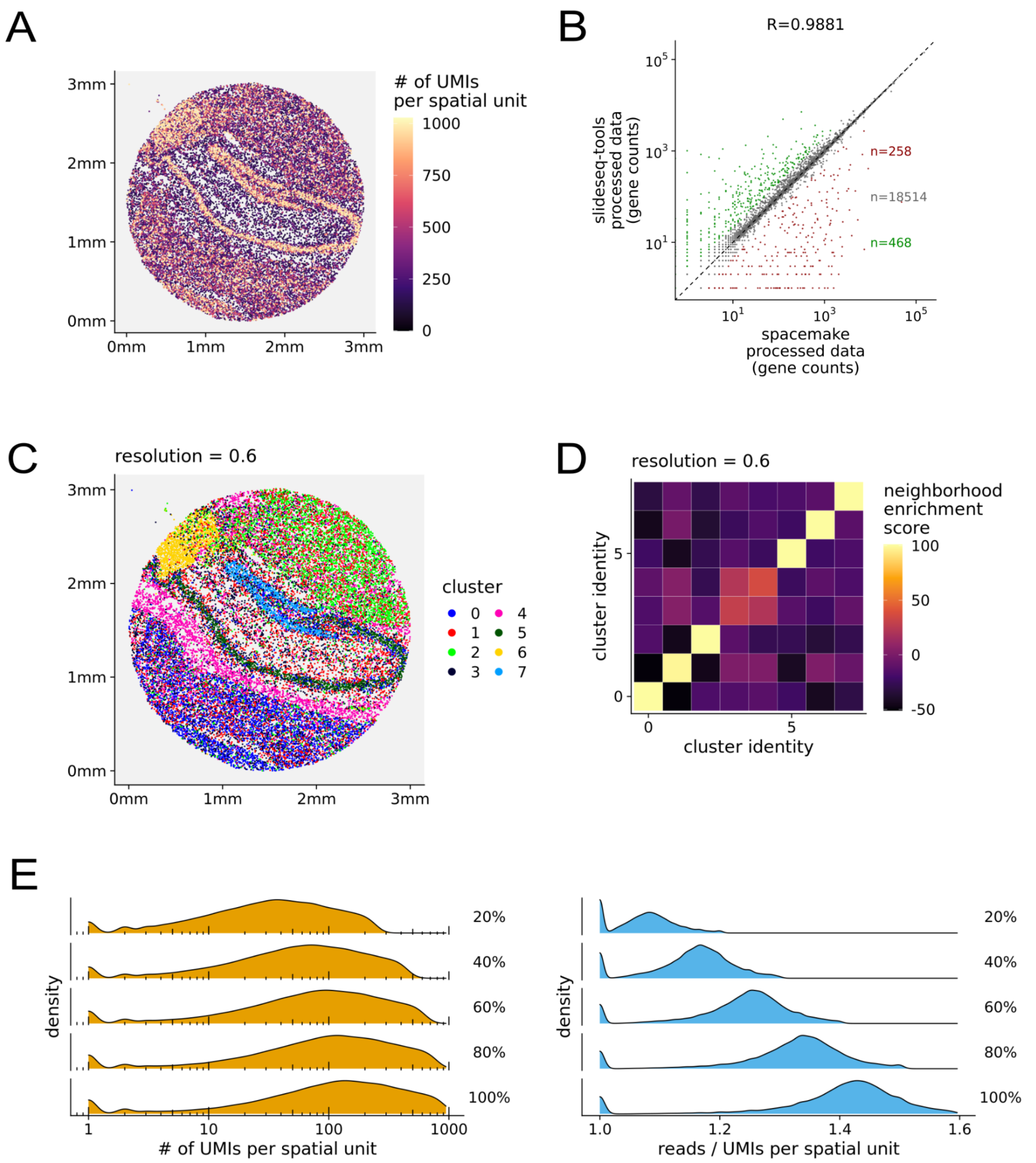


**Supplementary Figure 4.** Spacemake efficiently processes Slide-seqV2 data **(A)** Distribution of UMIs per bead over the puck. **(B)** Per-gene correlation of slideseq-tools and spacemake. Red genes are 2-fold enriched in slideseq-tools, while green genes are 2-fold enriched in spacemake. **(C)** Automated analysis identifies spatially resolved clusters, such as the cortical layer, dentate gyrus, pyramidal layer and thalamic region. **(D)** Neighborhood-enrichment with squidpy identifies clusters 3 and 4 to be neighboring in space. **(E)** Downsampling analysis reveals that distributions of UMIs per bead increase with sequencing depth (left) while the reads/UMIs ratio remains low (right), indicating that the sample has not reached sequencing saturation.

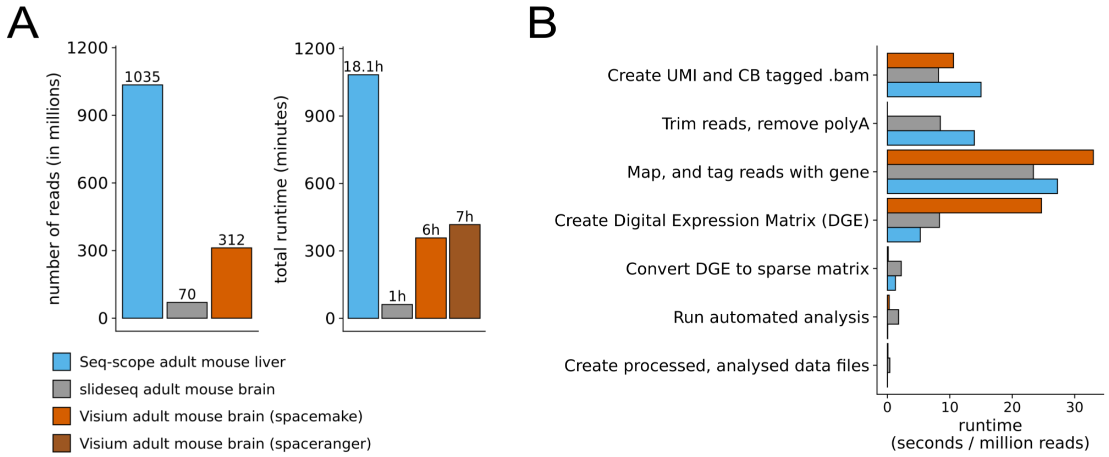


**Supplementary Figure 5.** Spacemake is fast, scales well and can simultaneously process multiple samples. **(A)** Spacemake is fast and is slightly faster than 10X spaceranger, while offering user-modifiable run-mode settings. Data here shown for a run-mode with spaceranger-like settings (multi-mapping reads counted, no poly(A) trimming, only exonic reads counted). **(B)** Spacemake scales well with increased library size. When normalized to the number of input reads, spacemake performs similarly regardless of sequencing depth.
